# CRISPR Screens Reveal Epstein-Barr Virus-activated JunB as a Key Lymphoblastoid B cell Dependency Factor that Represses Cyclin Dependent Kinase Inhibitor P18INK4c

**DOI:** 10.64898/2026.03.31.715635

**Authors:** Eric M. Burton, Yifei Liao, Davide Maestri, Bidisha Mitra, Benjamin E. Gewurz

## Abstract

Epstein-Barr virus (EBV) persistently infects over 95% of adults worldwide and is associated with a range of cancers, including lymphomas and epithelial malignancies. Despite advances in understanding EBV biology, targeted therapies for EBV-associated cancers remain limited. To identify novel dependencies in EBV-infected cancers, we performed genome-wide CRISPR-Cas9 loss-of-function screens in EBV+ lymphoblastoid versus Burkitt lymphoma cells, which differ by EBV latency programs. JunB emerged as a critical LCL-selective host dependency factor. LCL JunB knockout significantly decreased proliferation, with reduced G2/M progression, but without inducing apoptosis. JunB was more highly expressed in B cells with the EBV latency III than latency I program and correlated with LMPist1 levels in newly infected B cells. LMP1 stimulated JunB expression in a manner dependent on its cytoplasmic tail TES1/CTAR1 region and on canonical NF-κB. EBV-activated JunB played an obligatory role in repression of the G1/S phase inhibitor *CDKN2C*/p18^INK4c^ in LCLs but not Burkitt B cells. These findings establish an LMP1-JunB-p18^INK4c^ axis as essential for EBV-driven lymphoblastoid B cell proliferation, suggest JunB-mediated cross-talk between Epstein-Barr nuclear antigens and LMP1, and highlight JunB as a potential therapeutic target for EBV-associated lymphoproliferative disorders.

## Introduction

The gamma-herpesvirus Epstein-Barr virus (EBV) persistently infects >95% of adults worldwide. EBV causes 200,000 cancers per year, contributing to approximately 1% of total cancer burden. These cancers include endemic Burkitt lymphoma, Hodgkin lymphoma, post-transplant lymphoproliferative disease (PTLD) and HIV/AIDS-associated lymphomas[1, 2]. EBV also causes a range of epithelial cell tumors, including gastric and nasopharyngeal carcinomas (NPC), as well as T and NK cell lymphomas[3]. EBV is increasingly implicated as a key trigger for multiple autoimmune disorders, including multiple sclerosis[4–6] and systemic lupus erythematosus.

EBV colonizes the memory B cell compartment to establish the reservoir for lifelong infection. To accomplish this, EBV deploys discrete latency programs following B cell infection, in which combinations EBV latency genes are expressed. These include six Epstein-Barr nuclear antigens (EBNA), two latent membrane proteins (LMP) and non-coding RNAs[7–9]. The EBV genomic W promoter initially drives latency gene expression in newly infected cells. The fully transforming B-cell transforming latency III program is comprised of all six EBNA, LMP1 and LMP2A, and ncRNA and is driven by the EBV genomic C promoter. LMP1 and LMP2A mimic signaling from activated CD40 and B-cell receptors, respectively[10–12]. Latency III is hypothesized to drive infected B-cells into germinal centers, where infected B-cells switch to the latency IIa program. Latency IIa is comprised of EBNA1, LMP1 and LMP2A. The germinal center milieu may support LMP1 expression in the latency IIa program through JAK/STAT signaling[13–16]. Upon memory B cell differentiation, EBV switches to the latency I program, in which EBNA1 is the only EBV-encoded protein expressed[13, 17, 18].

LMP1 is expressed in many EBV-associated cancers, including PTLD, primary central nervous system lymphoma, Hodgkin lymphoma and nasopharyngeal carcinoma. LMP1 signaling is necessary for EBV-mediated human B cell immortalization into lymphoblastoid B cell lines. Transgenic LMP1 expression supports transformation of both B and epithelial cells[16, 19–24]. LMP1 is also implicated in multiple sclerosis pathogenesis[25–27].

LMP1 is comprised of a 24 residue N-terminal cytoplasmic tail, six transmembrane (TM) domains and a 200 residue C-terminal cytoplasmic tail[11, 28–30]. LMP1 TM domains drive ligand-independent, constitutive signaling from lipid raft sites[31–33]. Signaling from two LMP1 C-terminal tail regions, called transformation effector site (TES) or C-terminal activating region (CTAR) are each necessary for B-cell immortalization[34, 35]. TES1 activates canonical and non-canonical NF-κB, MAP kinase (MAPK), PI3 kinase, protein kinase C and JAK/STAT signaling. TES2 activates canonical NF-κB, MAP kinase, IRF7 and P62 pathways[11, 28, 30, 31, 35–42]. The LMP1 CTAR3 region activates JAK/STAT and SUMOylation pathways[43–45]. While canonical and non-canonical NF-κB signaling are each necessary for LCL growth and survival, considerably less is known about other LMP1-activated pathways that also support infected B cell proliferation.

Here, we built upon our prior CRISPR-Cas9 loss-of-function screen analysis[46] to identify host dependency factors important for EBV-transformed LCL or Burkitt B cell proliferation. Together with our prior screen that used a distinct CRISPR single guide RNA (sgRNA) library, this analysis highlighted key roles for the JunB AP-1 transcription factor in LCL growth. JunB knockout inhibited LCL but not Burkitt proliferation. We identified key LMP1 and JunB driven roles in suppression of the cell cycle G1/S phase inhibitor p18^INK4c^, encoded by *CDKN2C*.

## Results

### A human genome-wide CRISPR-Cas9 screen reveals lymphoblastoid B cell host dependency factors

To build upon our prior CRISPR analysis of host dependency factors important for EBV-transformed B-cell growth[46], we performed parallel CRISPR-Cas9 loss-of-function screens in the well characterized LCL GM12878 versus the EBV+ Burkitt lymphoma cell line P3HR-1. GM12878 is a tier I Encyclopedia of DNA Elements (ENCODE) cell line and expresses the viral latency III program. P3HR-1 expresses a more highly restricted EBV latency program, in which an EBV genomic deletion that knocked out EBNA2 drives triggered a Wp-restricted program[47–50], where EBNA1 is the major EBV latency program expressed, together with very low levels of other EBNAs and the EBV anti-apoptotic protein BHRF1. To build on our prior analysis with an earlier generation Avana sgRNA library which also used this cell line pair, we transduced Cas9+ GM12878 and P3HR1 with the Brunello sgRNA library[51]. Brunello is comprised of 77,441 lentiviruses that each encode one single-guide RNA (sgRNA, approximately 4 sgRNAs targeting each human gene). Brunello also contains ∼1000 non-targeting control sgRNAs[46]. Brunello has improved on-target and reduced off-target activity in comparison with the Avana library[52]. We used a multiplicity of infection of 0.3 to minimize lentivirus co-transduction. Successfully transduced cells were puromycin selected and passaged for 21 days, a format that has been widely used by the CRISPR DepMap project (DepMap.org). PCR-amplified sgRNA abundances were quantified via next-generation DNA sequencing. Hits, against which multiple sgRNAs were selectively depleted from GM12878 versus P3HR-1 and vice versa, were identified by the STARS algorithm[52] (**Fig. 1A, Supplementary Table 1**).

**Figure 1:**
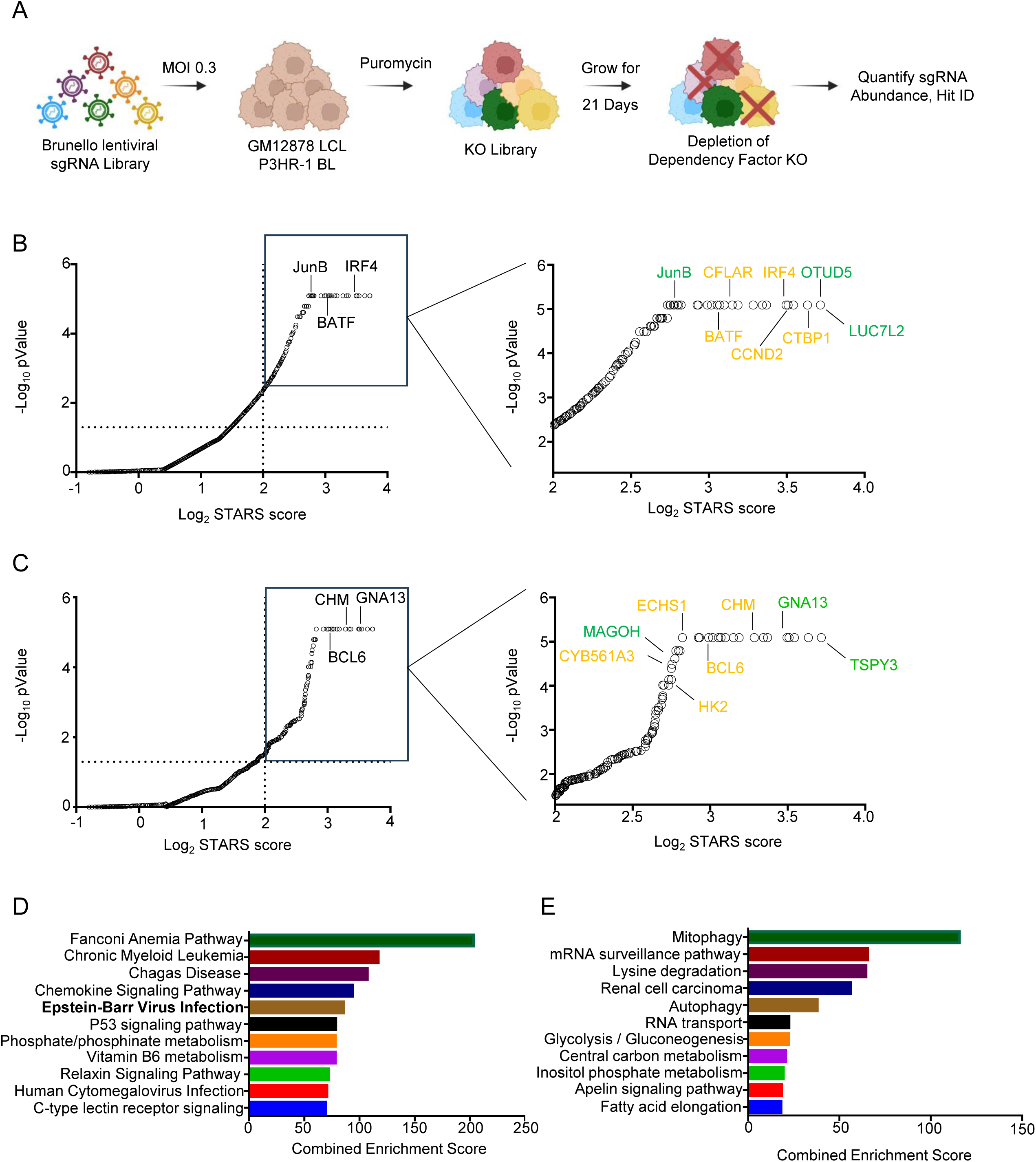
CRISPR-Cas9 screen for EBV-transformed B cell dependency factors. (A) CRISPR-Cas9 screen workflow. Cas9+ P3HR-1 Burkitt and GM12878 LCLs were transduced with the human genome-wide Brunello sgRNA library at a multiplicity of infection (MOI) of 0.3. Transduced cells were puromycin selected and passaged for 21 days. sgRNA abundances were determined via next generation sequencing of PCR-amplified genomic DNA. (B) Scatterplots of -Log10 (p-value) versus STARS algorithm scores. Shown at right is a zoomed in region highlighting LCL-selective hits, with hits that scored in this Brunello library screen but not in the prior Avana library screen green. GM12878-selective hits that scored in both the Avana and Brunello library screens are highlighted in yellow. Statistical significance was quantitated by the STARS algorithm, using two biological replicates for the Brunello screen. (C) Scatterplots of -Log10 (p-value) versus STARS algorithm scores. Shown at right is a zoomed in region highlighting P3HR1-selective hits, with hits that scored in this Brunello library screen but not in the prior Avana library screen green. P3HR-1-selective hits that scored in both the Avana and Brunello library screens are highlighted in yellow. Statistical significance was quantitated by the STARS algorithm, using two biological replicates for the Brunello screen. (D) KEGG pathway analysis of GM12878 LCL-selective Brunello screen hit dependency factors. (E) KEGG pathway analysis of P3HR-1 Burkitt-selective Brunello screen hit dependency factors.

At a multiple hypothesis test adjusted q<0.05 cutoff, we identified 158 hits whose sgRNAs were significantly more depleted from GM12878 versus P3HR-1 at the end of the screen. These were putative LCL-selective dependency factors, whose expression supported growth and/or survival of GM12878 LCL, but not of P3HR-1 Burkitt cells. Similarly, we identified 56 hits whose sgRNAs were selectively depleted in P3HR-1 versus GM12878 at Day 21 post-transduction and are therefore candidate host dependency factors for EBV+ Burkitt survival. Importantly, we identified multiple LCL selective hits that were in common with those identified in our published Avana sgRNA library screen[46], including the transcription factors IRF4, BATF, and CTBP1, CCND2 (which encodes cyclin D2) and CFLAR (which encodes the anti-apoptotic factor c-FLIP) (**Fig. 1B**, yellow). Similarly, multiple Burkitt-selective hits scored in our prior Avana library and current Brunello screen, including BCL6, CYB561A3[53], CHM, SLC16A1 and HK2 (**Fig. 1C**, yellow). LCL dependency factors were enriched for EBV infection, likely reflecting key roles of the EBV latency III oncogene-regulated host target genes (**Fig. 1D**). By contrast, we again found multiple metabolism pathways to be enriched amongst Burkitt-selective hits, including central carbon metabolism and glycolysis/gluconeogenesis, likely reflective of c-MYC driven oncometabolism (**Fig. 1E**).

### JunB is required for lymphoblastoid but not Burkitt B cell proliferation

LMP1 TES2 signaling activates MAPK signaling, and in the presence of TRAF1, LMP1 TES1 can also activate downstream MAPK pathways, including the c-Jun N-terminal kinase (JNK)[54]. LMP1 activated JNK phosphorylates Jun family members, which assemble into AP-1 transcription factors[55–58]. It was therefore notable that JunB was a top LCL-selective Brunello screen hit (**Fig. 1B**). Three of four Brunello *JunB* targeting sgRNAs were highly depleted in GM12878 following 21 days of culture, but not from P3HR-1 (**Fig. 2A**). Although JunB did not score as a hit in our prior Avana CRISPR screen, two of four sgRNAs were nonetheless highly depleted in GM12878 but not P3HR1 on day 21 (**Fig. S1A-B**). Interestingly, CRISPR analysis indicated that neither GM12878 nor P3HR-1 were dependent on Jun family members c-Jun or JunD for proliferation (**Fig. S1C-D**), indicating a specific LCL JunB dependency.

**Figure 2:**
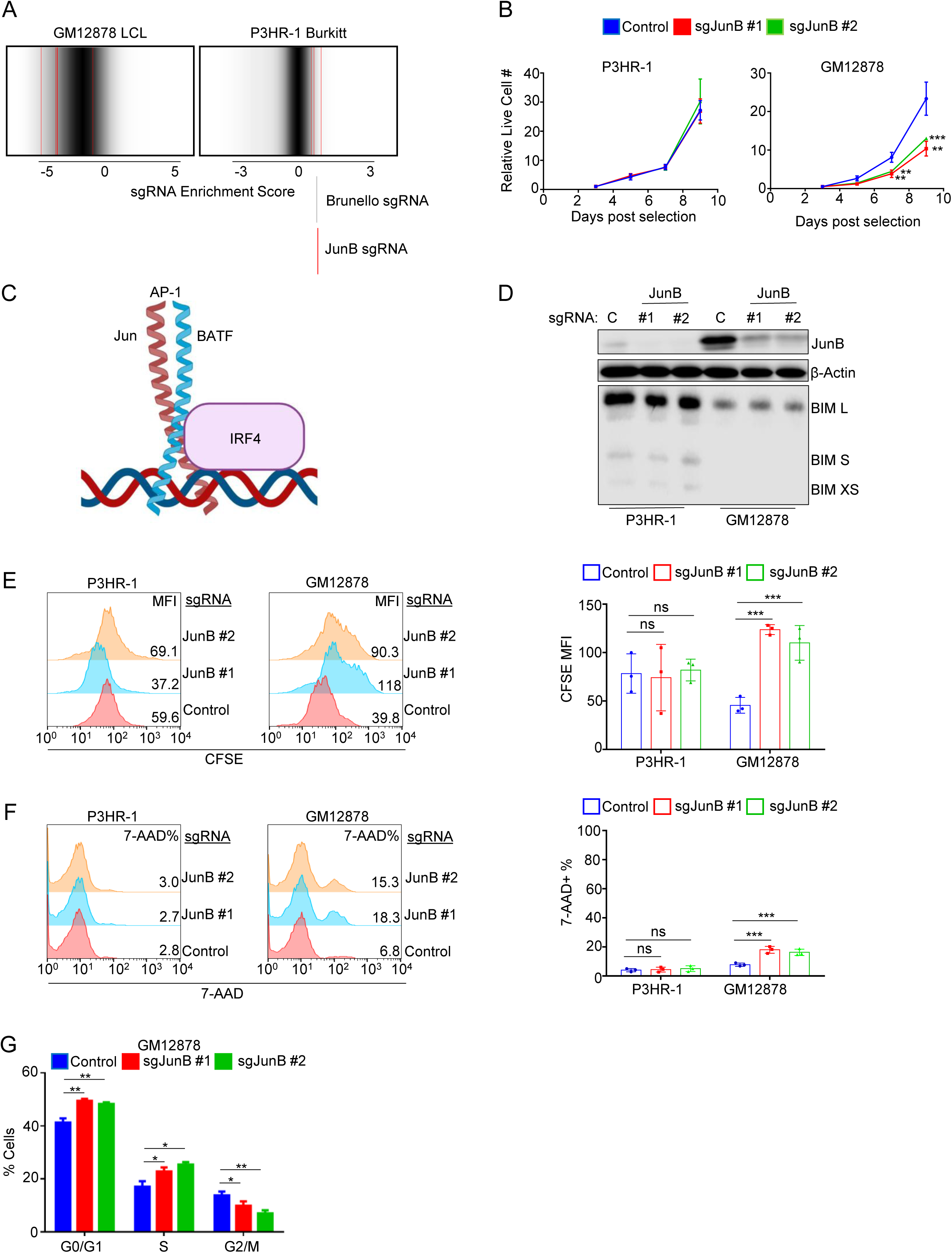
JunB is an LCL-selective dependency factor. (A) sgRNA enrichment score of the four guides targeting JunB (red) compared to the distribution of all sgRNA guides from the GM12878 and P3HR-1 Brunello dropout screens. A lower enrichment score indicates sgRNA depletion at day 21 versus day 1. Three of the four JunB targeting sgRNAs were highly depleted from the GM12878 library at day 21. All four were enriched in the P3HR-1 library at Day 21. (B) Mean ± SD live cell numbers from n=3 independent replicates of Cas9+ P3HR-1 (left) and GM12878 (right) expressing control or JunB targeting sgRNA. Cells transduced with lentiviruses expressing the indicated sgRNAs were puromycin selected. Cell numbers at two days following puromycin selection (defined as day 0) were set to 1. Live cell numbers were quantitated by CellTiter-Glo assay. (C) Jun-BATF-IRF4 complex schematic. AP-1 Jun/BATF transcription factors heterodimerize and can interact with IRF4 to regulate target genes. (D) JunB depletion does not increase BIM expression in P3HR01 or GM12878. Immunoblot analysis of WCL from Cas9+ P3HR-1 or GM12878 expressing control or JunB sgRNAs for nine days. Blots are representative of n=3 reps. (E) Analysis of JunB depletion effects on P3HR-1 and GM12878 proliferation. Cas9+ cells expressing control or JunB targeting sgRNA for 6 days were incubated for 96 hours following CFSE-staining and then analyzed by FACS. CFSE levels are reduced by half with each mitosis. Representative FACS plots are shown at left, while CSFE mean fluorescence intensity (MFI) ± SD from n=3 independent replicates are shown at right. (F) Analysis of Jun B depletion effects on P3HR-1 and GM12878 death. Cells from (E) were harvested ten days post-puromycin selection and stained with the vital dye 7-AAD, which is excluded from live cells. Representative FACS plots are shown at left, while mean ±%7-AAD+ values from n=3 experiments are shown at right. (G) Analysis of JunB depletion effects on P3HR-1 and GM12878 cell cycle. Cells as in (F) were fixed and stained with propidium iodide, and cell cycle stages were defined by FACS. Shown are mean ± SD values from n=3 independent replicates. P-values were determined by one-sided Fisher’s exact test. * p<0.05, **p<0.005, ***p<0.0005

We next inspected the DepMap[59] (Depmap.org) database of CRISPR-Cas9 dependencies across a library of cell lines. Notably, the proliferation of most DepMap cell lines is not dependent on JunB, with only 5 of 93 lymphoid cell lines found to be JunB dependent. Importantly, all five DepMap Burkitt cell lines were not dependent on JunB for proliferation (**Fig. S1E**). We validated that two independent screen hit Brunello JunB sgRNAs significantly reduced proliferation of GM12878 relative to non-targeting control values, but not P3HR-1 (**Fig. 2B**). Similar results were obtained in GM11830 LCLs versus Mutu I Burkitt cells (**Fig. S2A, B**), further validating JunB as an LCL-selective dependency factor. We also note that JunB levels were substantially higher in LCL than Burkitt whole cell extracts (**Fig. S2A**).

We previously reported that BATF and IRF4 are key LCL dependency factors[46], which form a complex that frequently co-occupies DNA sites with AP-1 transcription factors[60, 61] (**Fig. 2C**). We therefore hypothesized that JunB might be an LCL dependency factor based on its shared potential roles with BATF and IRF4. However, whereas a key BATF/IRF4 LCL role is to suppress apoptosis including by repressing BIM expression[46], JunB depletion by either of two sgRNAs failed to increase levels of BIM isoforms at day ten post sgRNA expression and puromycin selection, a timepoint when JunB depletion impaired proliferation (**Fig. 2D**). We therefore next investigated whether JunB depletion caused cell death or growth arrest. We performed parallel carboxyfluorescein succinimidyl ester (CFSE) dye-dilution and 7-AAD uptake assays on cells day 10 post control or JunB sgRNA expression, to read out effects on proliferation versus cell death, respectively. Whereas JunB depletion did not significantly reduce Burkitt proliferation or viability, it significantly decreased LCL proliferation, as judged by increased CSFE signal (CFSE is diluted by 50% with each mitosis) (**Fig. 2E-F**). By contrast, JunB depletion only modestly increased GM12878 cell death (**Fig. 2F**).

JunB can serve to antagonize c-Jun in certain contexts[62]. To investigate whether LCLs require JunB to counteract other Jun family members, we measured c-Jun and JunD levels in LCLs upon JunB depletion. However, we were unable to detect c-Jun expression, and JunB depletion did not increase JunD levels (**Fig S2C**). In agreement with our screen data, JunD KO did not perturb GM12878 growth, and combined JunB/D KO did not impair GM12878 proliferation more than JunB KO alone (**Fig S2D**). Taken together, this data suggests that LCL proliferation is dependent on JunB, independently of other Jun family members.

A key JunB role can be to regulate levels of cell cycle factors. It can downmodulate proliferation by increasing p16^INK4a^ (encoded by *CDKN2A*), while decreasing Cyclin D1 levels, yet in other cellular contexts JunB can instead promote proliferation, including by upregulating cyclin A2[63]. To therefore characterize potential JunB roles in LCL proliferation, we performed cell cycle analysis of GM12878 LCLs at day 10 post-sgRNA expression. JunB depletion increased the percentage of cells in G0/G1 and S phases but decreased the percentage of cells in G2/M (**Fig. 2G, S2E**).

### JunB is an EBV Latency III dependency factor

To gain further insights into the relationship between latency III and JunB, we first cross-compared JunB expression on the protein level across a panel of EBV+ B-cells with differing latency programs. JunB was more abundant in Mutu cells with the latency III than I program, and in latency III Jijoye than the P3HR-1 subclone that sustained an EBNA2 deletion and lost latency III expression (**Fig. 3A**). Likewise, JunB levels were generally higher in LCLs than in latency I Burkitt cells (**Fig. 3A**). To further assess this relationship, we utilized P493-6 LCLs, which have a conditional EBNA2 allele, whose nuclear localization is controlled by addition of 4-hydroxytamoxifen (4-HT). P493-6 also have a doxycycline repressible MYC allele[64]. Therefore, withdrawal of 4-HT and doxycycline converts the LCL into a Burkitt-like state with EBV latency I. Immunoblot analysis indicated that JunB levels were markedly higher when P493 grew in the latency III LCL state than in the latency I Burkitt-like state (**Fig. 3B**). Similarly, JunB levels progressively increased across the first 35 days of primary human peripheral blood B cell infection, where EBV converts resting cells into LCLs (**Fig. 3C**). Notably, levels of JunB correlated with LMP1 and with LMP1 target TRAF1 over this timecourse (**Fig. 3C**).

**Figure 3:**
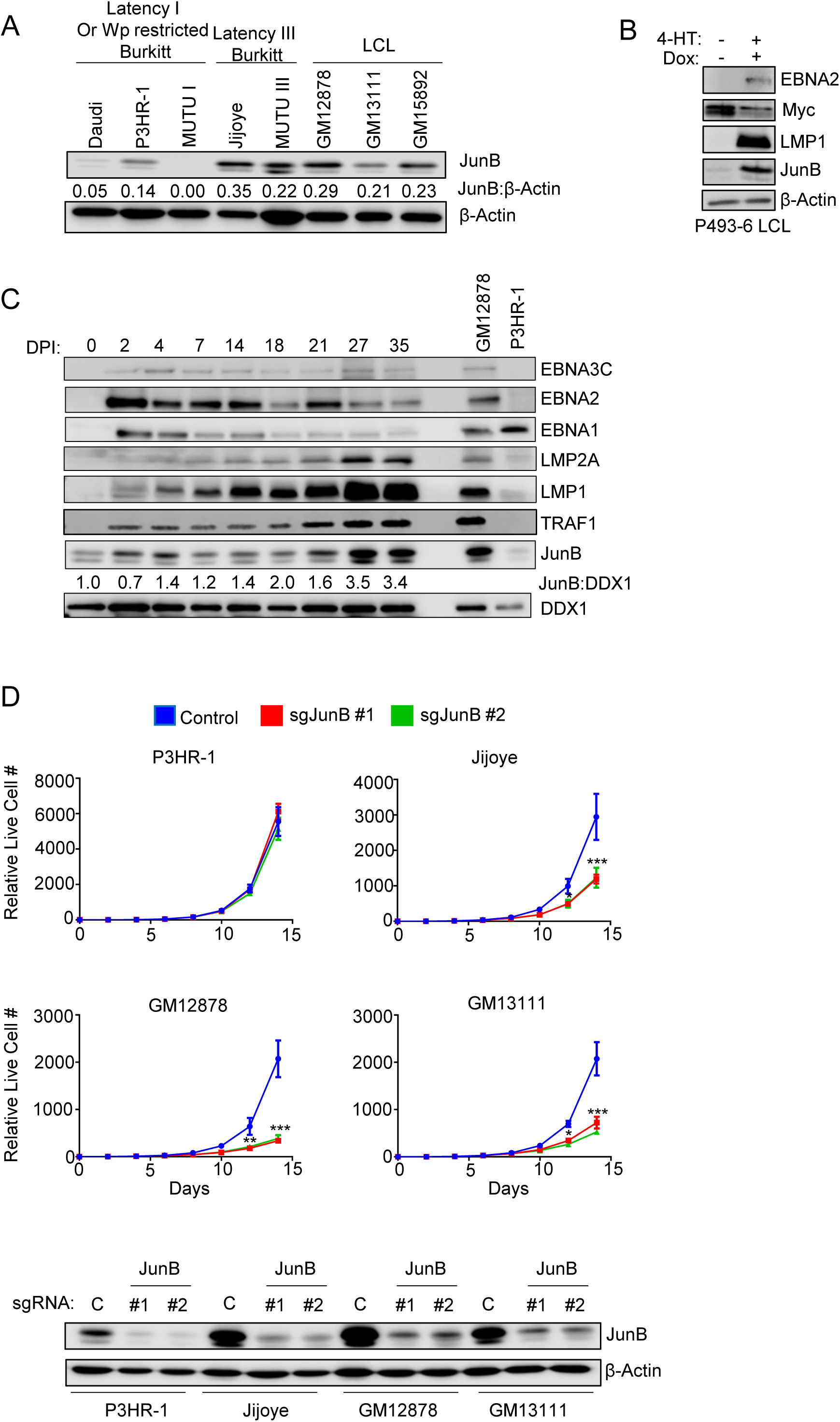
JunB is upregulated by and supports latency III B cell proliferation. (A) Analysis of JunB expression in B cells with latency III vs more highly restricted latency states. Immunoblot analysis of WCL from the indicated EBV+ B cells. (B) JunB expression in P493-6 cells grown in LCL vs latency I Burkitt-like states. P493-6 LCLs have a conditional EBNA2 allele, whose nuclear localization is dependent on the addition of 4-hydroxy tamoxifen (4-HT). P493-6 also have a tetracycline-repressed MYC allele. P493-6 were grown as latency III LCLs (+4-HT, +doxycycline) or in the Burkitt like state (−4-HT, -doxycycline) for 48 hours before WCL harvest and immunoblot analysis. (C) Analysis of JunB levels across EBV-mediated B cell immortalization. Peripheral blood B cells were infected with the EBV B95.8 strain. WCL was prepared from cells harvested at indicated days post-infection (DPI). JunB to DDX1 load control densitometry ratios are shown and were normalized to the Day 0 value. DDX1 was used as a load control as its levels do not vary substantially with EBV B cell transformation[75]. (D) Analysis of JunB depletion effects on latency III B cell proliferation. Shown are mean ± SD live cell numbers from n=3 replicates of Cas9+ P3HR-1, Jijoye, GM12878 or GM13111 that expressed control or JunB targeting sgRNA. Cell numbers two days following puromycin selection (defined as day 0 of the graph) were set to 1. Shown beneath are immunoblot analyses of WCL from cells with the indicated sgRNAs. P-values were determined by one-sided Fisher’s exact test. * p<0.05, **p<0.005, ***p<0.0005

To further test the relationship between latency III and JunB, we performed growth curve analysis on Jijoye versus P3HR-1 cells. Whereas JunB was dispensable for P3HR1 proliferation, in agreement with the CRISPR screen, it supported Jijoye proliferation (**Fig. 3D**). Even though Jijoye is a latency III Burkitt cell, its outgrowth was nearly as dependent on JunB as the LCLs GM12878 and GM13111 (**Fig. 3D**). Taken together, these data support a model in which latency III and potentially LMP1 induce JunB expression to support B cell outgrowth.

### LMP1 induces JunB expression

To build on the above results and prior reports implicating LMP1 in JunB upregulation[58, 65–67] we next functionally tested effects of LMP1 signaling on JunB expression. To build on microarray data that LMP1-driven NF-κB signaling upregulates JunB in IB4 LCLs[67], we tested effects of acute CRISPR LMP1 KO on JunB expression. In both GM12878 and GM13111 LCLs, LMP1 KO using a published sgRNA[68] rapidly depleted JunB (**Fig. 4A**). Since JunB mRNA has a half-life as short as 11 min[69, 70], these results are consistent with the need for LMP1 signaling to constantly maintain JunB levels in LCLs. In agreement with a key canonical NF-κB role, JunB upregulation in Daudi cells upon conditional doxycycline-induced LMP1 expression as blocked by small molecule IKKβ inhibition, in agreement with IB4 LCL results (**Fig. 4B**)[67].

**Figure 4:**
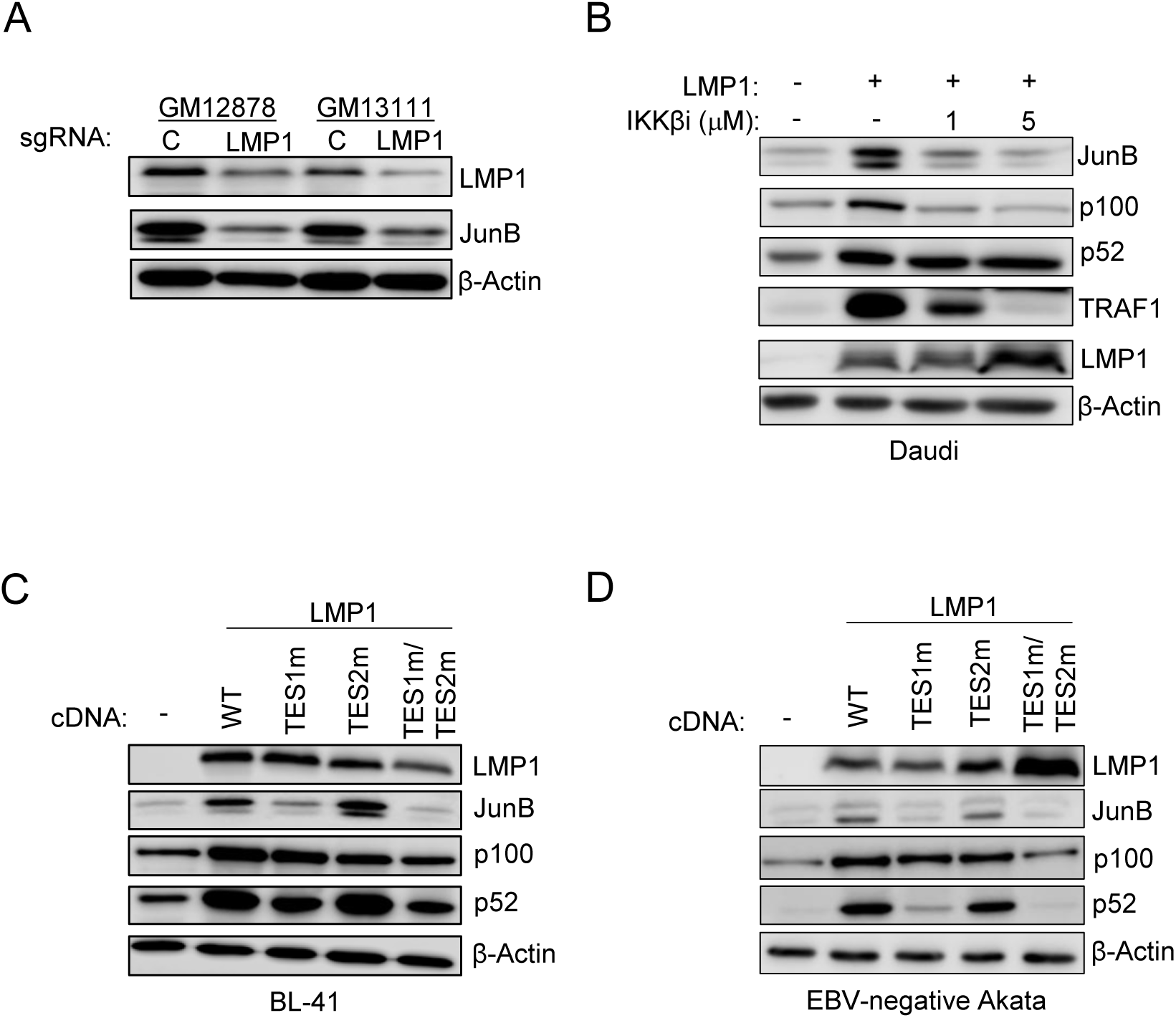
LMP1 TES1-mediated canonical NF-κB induces JunB expression. (A) LMP1 CRISPR KO effects on LCL JunB expression. Immunoblot analysis of WCL from Cas9+ GM12878 or GM13111 LCLs that expressed control or LMP1 targeting sgRNA, harvested three days post-sgRNA expression. (B) Analysis of role of NF-κB roles in LMP1-driven JunB induction. Immunoblot analysis of WCL from Daudi cells mock-induced or doxycycline (250ng/mL)-induced for conditional LMP1 expression for 24 hours and treated with DMSO vehicle or an IKKβ kinase inhibitor at 1 vs 5 μM, as indicated. (C) JunB induction by diverse NF-κB activating stimuli. Immunoblot analysis of WCL from Daudi cells that were mock stimulated or stimulated by multimerized CD40 ligand (CD40L, 50ng/mL), the SMAC mimetic birinapant (20μM) or TNFα (10ng/mL) for 24 hours. (D and E) Effects of LMP1 TES1 and TES2 signaling on JunB induction. Immunoblot analysis of WCL from BL-41 (D) or EBV-negative Akata (right) Burkitt cells that were mock-induced or doxycycline (250ng/mL) induced for wildtype (WT) or for transformation effector site (TES) 1 mutant (TES1m), TES2 mutant (TES2M) or TES1/TES2 double mutant (TES1m/TES2m) for 24 hours. Blots are representative of n=3 replicates.

We therefore next tested effects of conditional expression of wildtype LMP1 versus well characterized LMP1 point mutants abrogated for signaling from TES1 (TES1 mutant, TES1m), from TES2 (TES2m) or both (TES1m/TES2m)[71] [71–73]. Wildtype and TES2 mutant LMP1 robustly induced JunB expression in two EBV-negative Burkitt cell models, BL41 and Akata (**Fig. 4C-D**), whereas LMP1 TES1m or TES1m/TES2m failed to do so (**Fig. 4C-D**). These data are therefore consistent with a model in which an LMP1 TES1 driven canonical NF-κB pathway induces JunB expression.

### JunB inhibits p18^INK4c^ to promote B cell proliferation

Consistent with results from our Avana sgRNA library CRISPR screen[46], sgRNAs targeting CCND2, which encodes cyclin D2, were highly depleted from GM12878 but not P3HR1 (**Fig. S3A**). Notably, EBV also progressively induced cyclin D2 RNA (**Fig. S3B**) and protein[74, 75], with similar kinetics as LMP1 and JunB induction. We therefore tested whether conditional LMP1 expression was sufficient to induce cyclin D2 in latency I Daudi cells. Interestingly, WT LMP1 strongly induced cyclin D2, to a much greater extent than TES1m or TES2m LMP1 expression (**Fig. S3C**). However, JunB knockout did not significantly alter cyclin D2 expression in either Jijoye or GM12878 cells (**Fig S3D**).

To therefore gain insights into how JunB instead supports latency III B cell proliferation, we performed RNAseq on Cas9+ GM12878 following expression of control versus JunB sgRNAs using n=3 replicates. Interestingly, of the 307 differentially expressed genes at a 2-fold cutoff, mRNAs encoding the T-cell attractive chemokines CXCL9 and CXCL10 were amongst the most highly JunB KO upregulated transcripts (**Fig. 5A, Supplementary Table 2**). Notably, EBNA3A negatively regulates each through polycomb repressive complex two dependent histone 3 lysine 27 trimethylation (H3K27me3)[76, 77], suggesting cross-talk between LMP1, JunB and ENBNA3A. *CDKN2C* mRNA, which encodes the cyclin dependent kinase inhibitor p18^INK4c^, was also highly upregulated by LCL JunB depletion (**Fig. 5A**, **Table 1**). No other CDKN2C isoform was differentially expressed due to JunB depletion.

**Figure 5:**
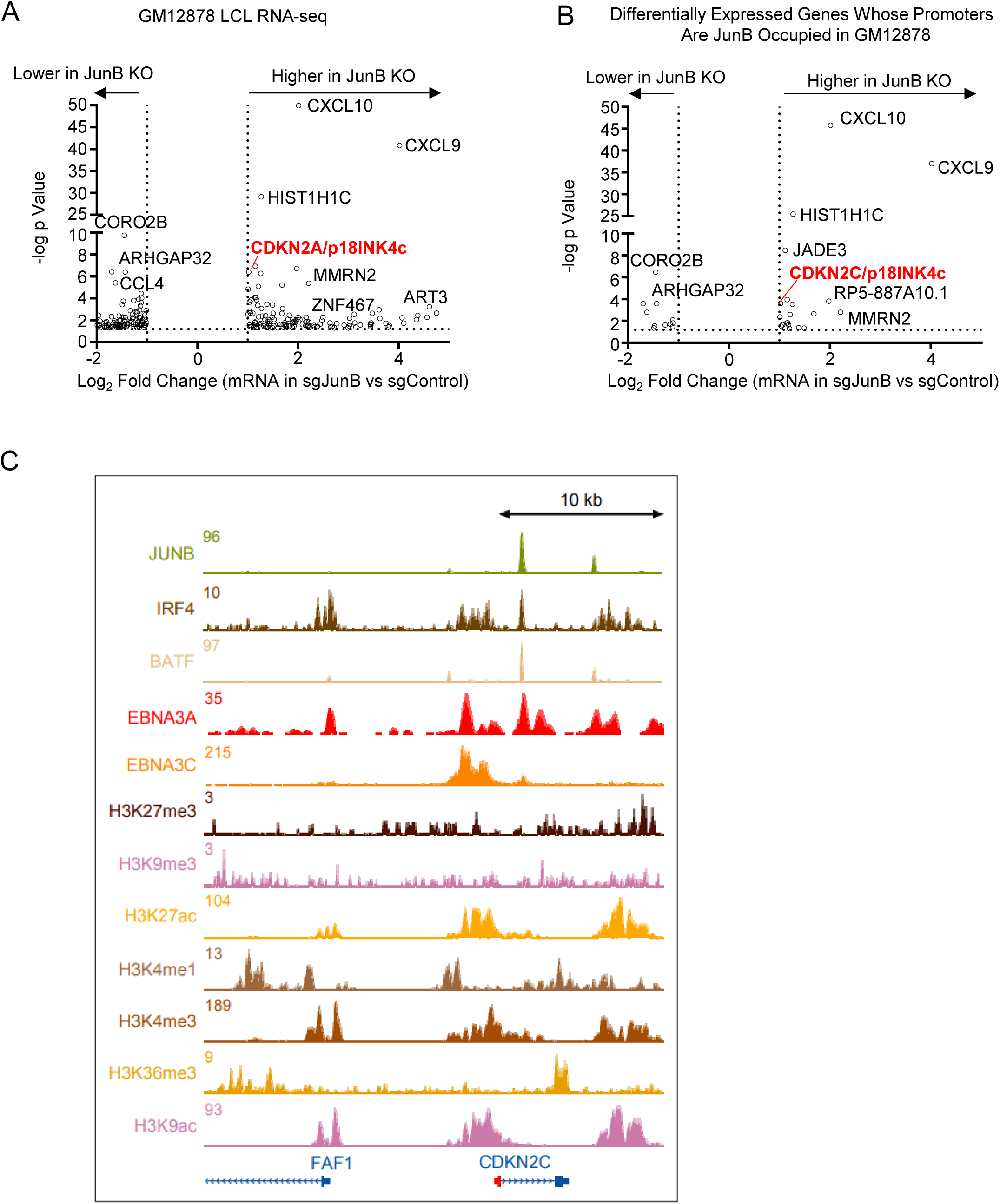
Analysis of JunB roles in LCL target gene regulation. (A) RNAseq volcano plot analysis of mRNAs with significantly different expression in control versus JunB KO LCLs. Triplicate RNAseq analyses were performed on Cas9+ GM12878 LCLs with control or JunB sgRNA. RNAseq was performed ten days post-puromycin selection. Transcripts whose abundance were Log2 >1 or <1 different in JunB KO cells are shown. (B) Analysis of LCL JunB target genes. RNAseq volcano plot analysis as in (A), showing expression values for differentially expressed mRNAs for whose promoters were also bound by JunB within 1 kb of transcription start sites, using GM12878 ENCODE JunB ChIP-seq data[132]. (C) LCL ChIP-seq tracks at the CDKN2C locus. Shown are normalized ChIP-seq signals for JunB, EBNA3/C, IRF4, BATF, or for the indicated histone methylation or acetylation marks.

**Table 1.**
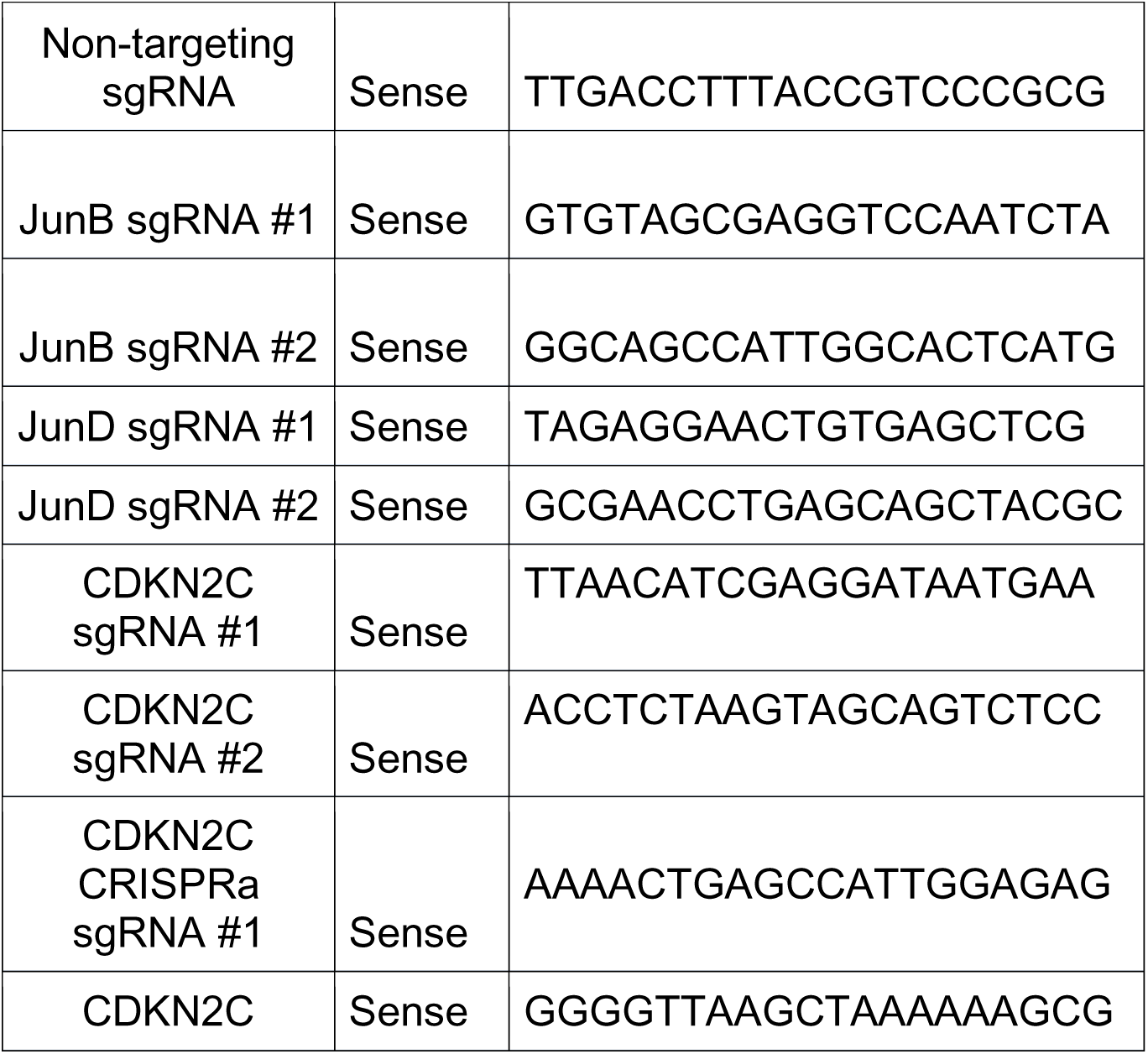
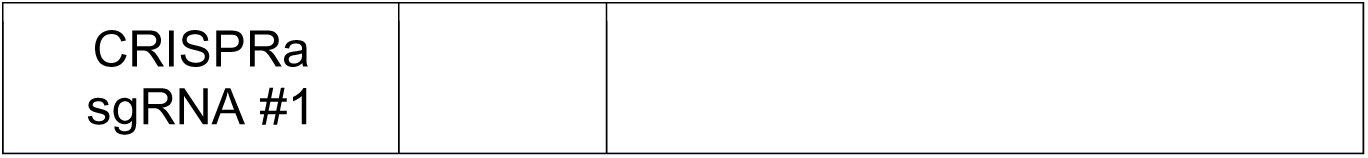
CRISPR-Cas9 and CRISPRa sgRNA sequences.

We next cross-compared our RNAseq with JunB ChIP-seq analysis performed in GM12878 by the Encyclopedia of DNA Elements (ENCODE) project[78, 79]. Altogether, 30 differentially regulated genes had JunB occupancy within 1kb of their promoters, suggesting that most JunB differentially regulated genes are likely to be indirectly regulated by JunB or regulated at distal regulatory elements (**Fig. 5B**). This analysis highlighted that *CDKN2C* mRNA was also amongst the most highly upregulated transcript of LCL JunB bound genes, for whom JunB peaks were within 1kb of the transcription start site (**Fig. 5B**). JunB, IRF4 and BATF co-occupied a site downstream of the *CDKN2C* promoter. LCL EBNA3A and 3C ChIP-seq datasets[76, 80] further highlighted that EBNA3A also co-occupied this site (**Fig. 5C**), whereas EBNA3C highly occupied a nearby site just upstream of the *CDKN2C* promoter.

ENCODE project GM12878 ChIP-seq epigenetic analyses further highlighted the presence of activating histone 3 lysine 27 acetyl, histone 3 lysine 9 acetyl, histone 3 lysine 4 monomethyl and trimethyl marks near the *CDKN2C* promoter, with relatively low level of repressive histone 3 lysine 27 and histone 3 lysine 9 trimethyl marks(**Fig. 5C**). Taken together, this suggests that the LCL *CDKN2C* locus is epigenetically poised for transcription and support a model in which JunB co-occupies the LCL *CDKN2C* locus together with BATF, IRF4 to recruit EBNA3 proteins and co-repressors to block *CDKN2C*/p18^INK4c^ expression.

### JunB repression of LCL proliferation is partially dependent on p18INK4c

*CDKN2C* encoded p18^INK4c^ inhibits cyclin dependent kinases 4 and 6 (CDK4/6) to trigger G1 cell cycle arrest[81, 82]. Interestingly, p18INK4c expression is epigenetically repressed by EBNA3A and 3C, in part through polycomb repressive complex II [83]. To gain insights into functional p18INK4c roles in Burkitt vs LCL contexts, we used CRISPR activation (CRISPR-a)[84] to cross-compare effects of p18^INK4c^ upregulation in these two EBV+ B cell contexts. CRISPR-a strongly induced p18^INK4c^ expression in P3HR-1 and in GM12878 (**Fig. 6A**). To define *CDKN2C* CRISPR-a effects on P3HR-1 vs GM12878 proliferation, live cell numbers were quantitated after 7 days of control versus *CDKN2C*-targeting sgRNA expression. Importantly, despite higher p18^INK4c^ levels in P3HR-1, p18^INK4c^ over-expression more strongly repressed GM12878 than P3HR-1 proliferation (**Fig. 6B**). Similar results were obtained in a second Burkit/LCL pair, using MUTU I vs GM15892 cells (**Fig. 6C-D**).

**Figure 6.**
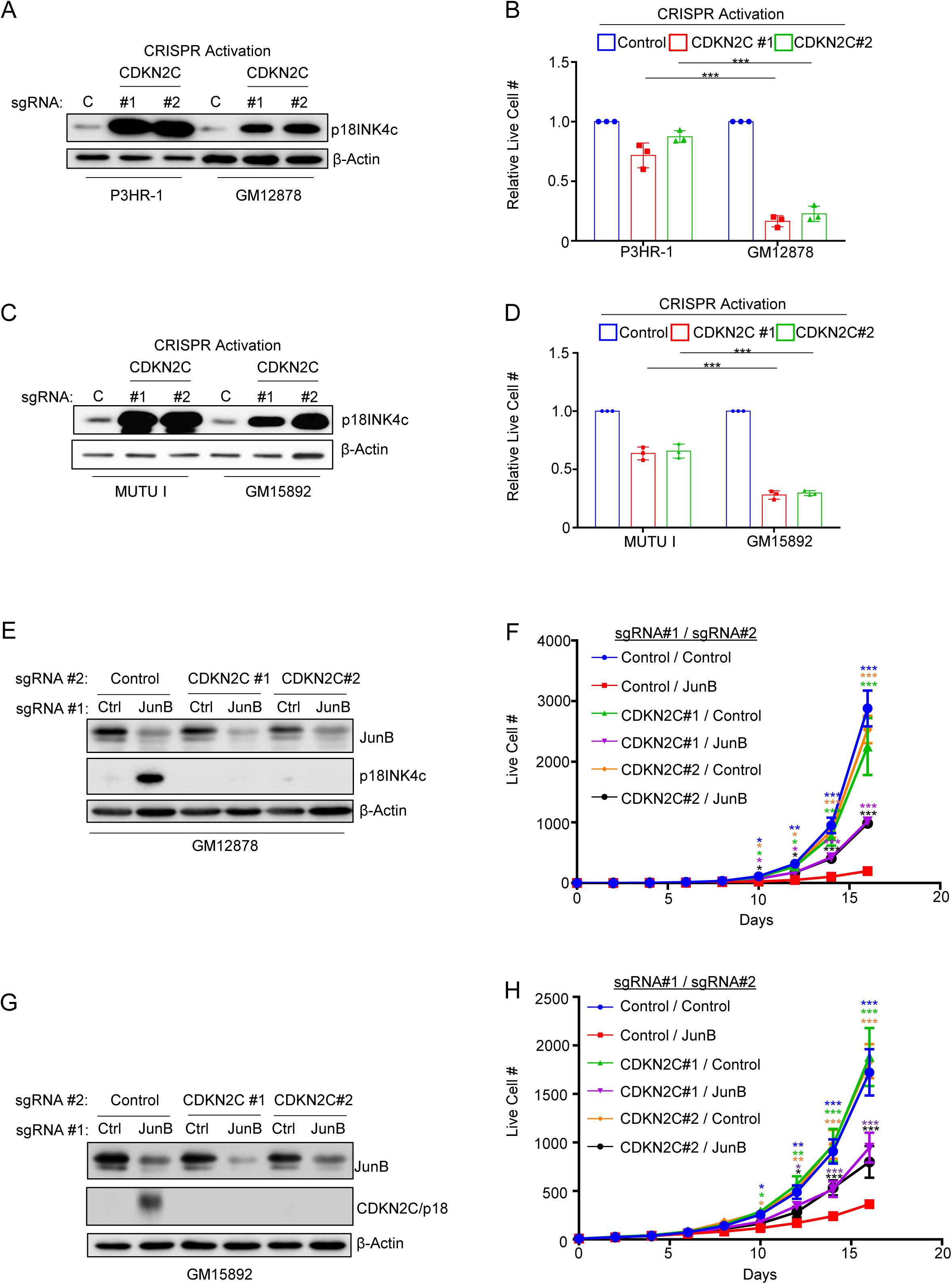
JunB represses CDKN2C/P18INK4c to support LCL proliferation. (A) CRISPR activation (CRISPR-a) overexpression of p18INK4c in P3HR-1 Burkitt and GM12878 LCL. Immunoblot analysis of WCL from CRISPR-a (dCas9-VP64+) P3HR-1 and GM12878 cells that expressed control or CDKN2C targeting sgRNAs. (B) Effects of p18IKF4c over-expression on P3HR-1 versus GM12878 proliferation. Mean ± SD live cell numbers of cells as in (A) from n=3 replicates. Cells were re-seeded four days following transduction and puromycin selection. Live cell numbers were quantitated on day seven post-puromycin selection. (C) CRISPR-a overexpression of p18INK4c in Mutu I Burkitt and GM15892 LCL. Immunoblot analysis of WCL from dCas9-VP64+ Mutu I and GM15892 cells that expressed control or CDKN2C targeting sgRNAs. (D) Effects of p18IKF4c over-expression on Mutu I versus GM15892 proliferation. Mean ± SD live cell numbers of cells as in (C) from n=3 replicates. Cells were re-seeded four days following transduction and puromycin selection. Live cell numbers were quantitated on day seven post-puromycin selection. (E) Combined LCL JunB and CDKN2C KO. Immunoblot analysis of WCL from Cas9+ GM12878 that expressed the indicated control or JunB sgRNAs and that also expressed the indicated control or CDKN2C targeting sgRNAs. (F) CRISPR genetic analysis of CDKN2C roles in GM12878 LCL growth arrest upon JunB KO. Growth curve analysis of Cas9+ GM12878 LCLs that expressed the indicated sgRNAs as in (E). (G) Combined LCL JunB and CDKN2C KO. Immunoblot analysis of WCL from Cas9+ GM15892 that expressed the indicated control or JunB sgRNAs and that also expressed the indicated control or CDKN2C targeting sgRNAs. (H) CRISPR genetic analysis of CDKN2C roles in GM15892 LCL growth arrest upon JunB KO. Growth curve analysis of Cas9+ GM15892 LCLs that expressed the indicated sgRNAs as in (E). Blots are representative of n=3 replicates. one-sided Fisher’s exact test * p<0.05, **p<0.005, ***p<0.0005

We next asked to what extent LCL proliferation was dependent upon *CDKN2C* repression by JunB. To do so, we first expressed control or independent CKDN2C targeting sgRNAs in Cas9+ GM12878 or GM15892 LCLs. We then expressed control or JunB targeting sgRNAs. Successful CRISPR knockout was validated by immunoblot, which showed appropriate JunB depletion as well as p18^INK4c^ upregulation upon JunB KO in control, but not in *CDKN2C* edited cells (**Fig. 6E-F**). We then performed growth curve analysis to define effects of JunB KO, alone or in the presence of combined p18^INK4c^ KO. Interestingly, whereas JunB KO very strongly suppressed proliferation of both LCLs, combined p18^INK4c^ depletion partially rescued their outgrowth (**Fig. 6G-H**). Collectively, our results support a model in which LMP1 upregulates JunB, which together with EBNA3A/C represses p18^INK4c^ to support LCL proliferation (**Fig. 7**).

**Figure 7:**
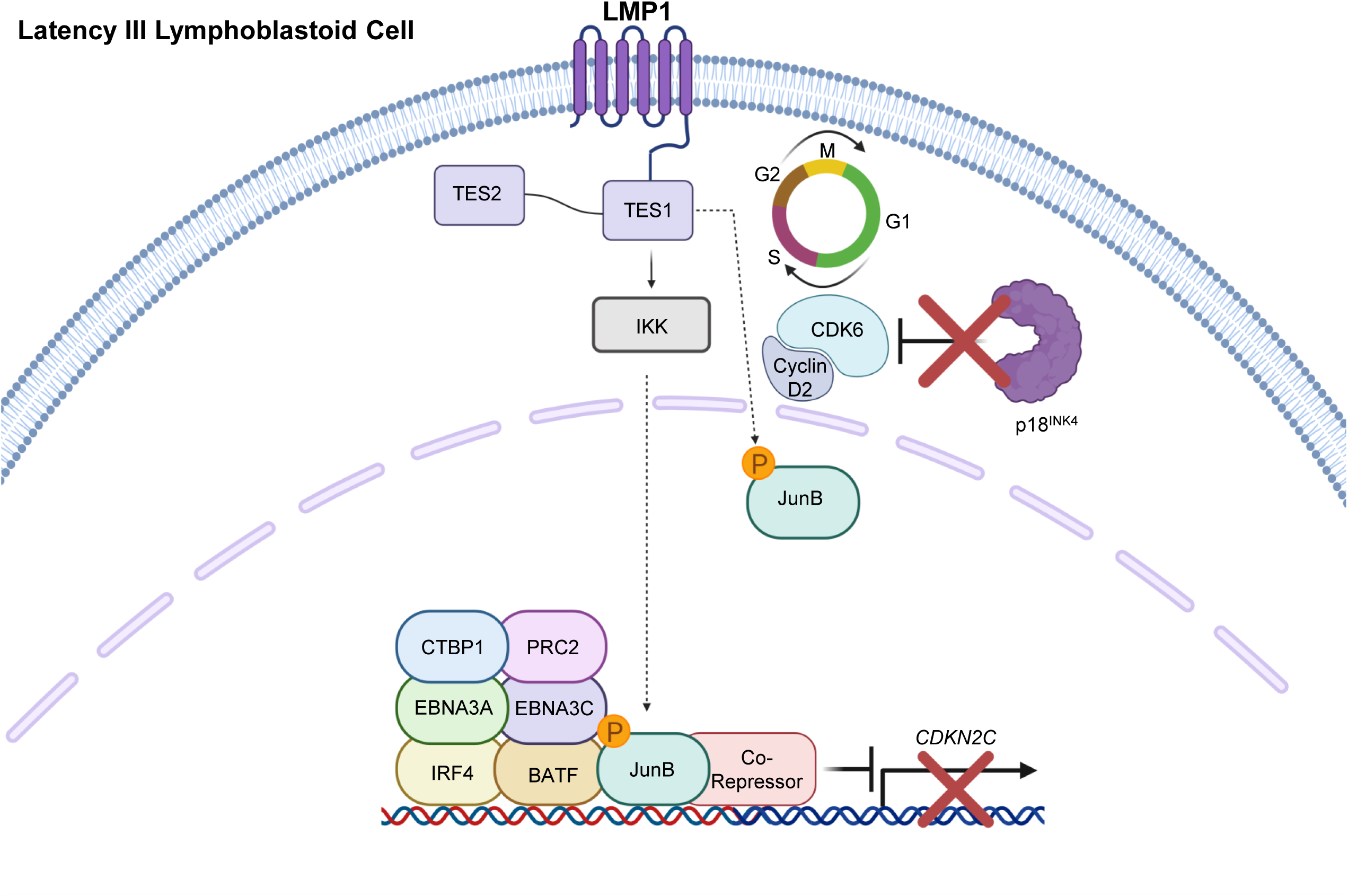
Model of JunB cell cycle regulation via CDKN2C/p18INK4c silencing. LMP1 TES1 signaling induces JunB and drives its nuclear translocation. A key JunB role is to repress CDKN2C expression, likely together with BATF/IRF4, EBNA3A, and 3C. In the absence of JunB, CDKN2C-encoded P18INK4c is expressed and blocks CDK6/Cyclin D2 to impair cell cycle progression. Created in BioRender. Burton, E. (2026) https://BioRender.com/gr5ur3n

## Discussion

Here, we used parallel human genome-wide CRISPR-Cas9 screens to identify that JunB is an LCL-selective dependency factor. JunB KO impaired LCL, but not Burkitt B cell proliferation. The EBV latency III program induced JunB expression, which was found to be driven by LMP1 TES1 and canonical NF-κB signaling. LCL JunB KO de-repressed expression of CDKN2C encoded 18^INK4c^, which inhibits another LCL dependency factor, CDK6. By contrast, p18^INK4c^ over-expression strongly blocked LCL but not Burkitt B cell proliferation. Furthermore, CRISPR p18^INK4c^ depletion partially rescued LCL growth upon JunB KO. Taken together, our results highlight CDKN2C as a key LMP1 and JunB target.

Upon primary B cell infection, EBNA3A and 3C epigenetically repress *CKDN2C* within the first 12 days post-infection[83]. However, if they are conditionally inactivated after this timepoint, *CDKN2C* remains repressed in a manner that is partially dependent on polycomb repressive complex 2 (PRC2) activity[83]. By contrast, p18^INK4c^ levels markedly increase at days 15-20 post-infection in B cells infected by EBNA3A/C knockout EBV[83]. Interestingly however, LCL ChIP-seq demonstrates only modest *CDKN2C* locus deposition of PRC2 histone 3 lysine 27 trimethylation (H3K27me3) repressive marks. While combined EBNA3A/C inactivation fails to de-repress p18^INK4a^ LCL expression, JunB KO was sufficient to de-repress its LCL expression.

Our data therefore support a model in which EBNA3A/C and LMP1 cross-talk to repress *CKDN2C/*p18^INK4a^. Notably, JunB co-occupies a LCL *CKDN2C* locus site together with BATF and IRF4, suggesting that it may be present as a heterodimer with BATF that interacts with DNA-bound IRF4. ChIP-seq further highlights that EBNA3A and EBNA3C often co-occupy host genomic sites with BATF and IRF4[46, 76, 85], and IRF4 is enriched at both EBNA3A- and EBNA3C-bound LCL sites[86]. IRF4 is necessary for EBNA3C binding to specific sites[86]. We anticipate that JunB nucleates repressor complexes that include EBNA3A, EBNA3C and additional host co-repressor(s) to block LCL *CDKN2C/*p18^INK4a^ expression. In support, JunB has established roles in gene repression in the contexts of Th17 differentiation[87], in repression of the gene encoding G-CSF in keratinocytes[88, 89], and in TGF-β repression[90]. Therefore, an important future objective will be to identify nuclear co-repressors recruited in a JunB-dependent manner to the LCL *CDKN2C* locus.

The CDK inhibitor family includes p16^INK4a^, p15^INK4b^ and p18^INK4c^. P18^INK4c^ has key B-cell specific roles, where it terminates cell proliferation and promotes plasma cell differentiation[91–93]. Mice deficient in p18^INK4c^ exhibit B cell hyperproliferation, both within germinal centers and within extrafollicular foci[91]. Due to the major defect in plasma cell differentiation, they have severely reduced antibody titers. The major p18^INK4a^ B cell target is CDK6[93]. p18^INK4a^ blocks CDK6 interaction with D-type cyclins, thereby impairing phosphorylation of Rb to trigger G1/S phase cell cycle arrest. This activity supports terminal B cell differentiation[82, 93]. Our results also raise the question of whether LMP1-driven JunB expression also silence *CKDN2C* in cells with the latency II program (EBNA1/LMP1/LMP2A), including germinal center B-cells or Hodgkin lymphoma Reed-Sternberg cells. Limited sequencing data is currently available for either of these cell states, though single cell approaches should increasingly allow their characterization. JunB is highly expressed in at least a subset of Hodgkin tumors[94–97], though EBV effects remain to be studied in this context.

LMP1 activates JNK[12, 24, 34, 38, 40, 54–57, 98–106], which phosphorylates Jun transcription factors. Interestingly, we observed greater LMP1-mediated JunB upregulation on the protein level. An interesting area of future investigation will therefore be to determine wither LMP1 signaling increases JunB stability, potentially through post-translational modification. It will also be of interest to define whether LMP1 and JunB repress *CDKN2C* in the context of nasopharyngeal carcinoma to support EBV-transformed cell growth. Interestingly, LMP1 activates JunB and supports nasopharyngeal carcinoma proliferation[65], and elevated JunB expression correlates with poor NPC outcomes[107]. LMP1 also downregulates CDKN2A/p16^INK4a^ in epithelial cells by promoting E2F4/5 nuclear export[108]. Additionally, LMP1 induces the inhibitor of differentiation (Id) 1 and 3 proteins in epithelial cells, which in turn suppress CDK2NA and CDKN1B to promote CDK4/6 and CDK2 activity[109].

EBNA3C and to a lesser extent EBNA3A repress CDK2NA/p16^INK4a^ to support EBV-driven B-cell immortalization *in vitro* and in murine lymphoma models[110–117]. Additionally, EBNA3A represses the cyclin-dependent kinase inhibitor p21/WAF1/CIP1[118]. LMP1 induces cyclin D2 expression[119], and LMP1 over-expression overrides serum starvation-induced growth arrest[120]. Taken together with our finding that CDK6 and cyclin D2 are LCL-selective dependency factors[46], our data suggest that LMP1 evolved multiple roles to support cell cycle. Therefore, latency III evolved a wide range of non-redundant mechanisms to highly remodel the cyclin-dependent kinase inhibitor landscape in support of aberrant B cell proliferation.

EBV oncoproteins are critical for lymphoblastoid B cell growth but have not yet proven to be druggable. Our results highlight multiple host factors downstream of EBV oncoprotein that are potential therapeutic targets for blockade of EBV-driven B cell oncogenic growth. These include CDK6, for which FDA-approved drugs are in clinical use, including palbociclib, as well as the LMP1/JunB pathway. Since inhibition of the polycomb repressive complex II EZH2 enzymatic subunit moderately interfered with EBNA3A/C repression of p18^INK4^[83], it will be of interest to test whether palbociclib and EZH2 inhibitors in clinical development additively or synergistically induce lymphoblastoid B cell p18^INK4c^. Likewise, we suspect that JNK1/2 did not score in our CRISPR screen due to functional redundancy, as the JNK inhibitor SP600125 blocks LCL growth *in vitro* and in murine xenografts[98]. It will therefore be of interest to test whether JNK inhibitors in clinical development exhibit additively or synergistically retrain LCL proliferation together with EZH2 or CDK6 inhibitors.

In summary, our CRISPR-Cas9 screen identified JunB as a key LCL-selective dependency factor that supports growth of EBV-infected B cells with the transforming latency III program. LMP1 TES1 signaling upregulated JunB, which was required for repression of the cell cycle inhibitor p18^INK4c^. Since the key p18^INK4c^ target CDK6 is also an LCL dependency factor, our study suggests that LMP1-driven JunB activity is required to block p18^INK4c^, together with EBNA3A/C, to support CDK6-driven proliferation. These results highlight therapeutic targets for the inhibition of EBV lymphoblastoid B-cell proliferation, including in post-transplant lymphomas.

## Acknowledgements

EMB and BM were supported by American Cancer Society Post-doctoral Fellowships PF-23-898493-01-TBE and PF-24-1250090-01-IBCD, respectively. These studies were also supported by R01CA228700, R01DE033908, P01CA269043 and U01CA275301 to BEG, and by a philanthropic support from George and Sandi Schussel to BEG. We thank Jaap Middeldorp for generously providing the OT1X anti-EBNA1 monoclonal antibody. CRISPR screen and RNAseq datasets have been deposited in the NIH GEO omnibus and will be released upon publication.

## Materials and Methods

### Ethics statement

Platelet-depleted venous blood obtained from the Brigham & Women’s hospital blood bank were used for primary human B cell isolation, following our Institutional Review Board-approved protocol for discarded and de-identified samples. For use in RNA-seq, Platelet-depleted venous blood was purchased from Gulf Coast Blood Center. The Mass General Brigham Hospital Institutional Review Board (IRB) approved this study, protocol 2022P001270 and 2019p002995. Formal consent was obtained by the Brigham & Women’s hospital blood bank during before donation.

### Pooled genome-wide CRISPR screens

A total of 150 million Cas9^+^ GM12878 or P3HR-1 cells were spin-inoculated for 2 h at 300*g* in the presence of 4 μg ml^−1^ polybrene in 12-well tissue culture plates, with the Brunello sgRNA library[51] at a multiplicity of infection (MOI) of ∼0.3. Brunello targets each human gene with four independent sgRNAs on average and includes 1,000 non-targeting negative controls. The plates were incubated at 37 °C for 6 h after spin-inoculation and then were cultured in fresh growth medium at 0.3 million per milliliter density. The puromycin (3 μg ml^−1^) was added at 48 h post transduction. Biological duplicate libraries were then passaged every 72 hours, returning cell number to at least 40 million per library at each passage to maintain adequate complexity. After 21 days of passaging, Genomic DNA was extracted using the QIAGEN Blood and Cell Culture DNA Maxi Kit (Qiagen, no. 13362). All DNA were sent for PCR amplification and sequencing to quantify the sgRNA abundance. The statistically significant screen hits were identified following the STARS algorithm analysis[52]. A multiple test hypothesis adjusted q-value < 0.05 from the STARS analysis was defined as a significant hit. sgRNA enrichment (as in Fig. 2A, S1A) scores were visualized in ggplot2. The output was displayed in two layers tiled across a one-dimensional distribution. The first layer was calculated by computing a kernel density estimate (https://ggplot2.tidyverse.org/reference/geom_density.html) over all input values. The density values are represented by fill color, with darker regions having more density and lighter regions having less density represent individual sgRNA abundance. The second layer plotted a subset of individual values belonging to the gene of interest. Values for gene of interest were indicated in red.

### Cell lines and culture

293T, Daudi and Jijoye were purchased from American Type Culture Collection. P3HR-1, Daudi, and EBV-negative Akata[121] were obtained from Elliott Kieff. GM11830, GM12878, GM13111, and GM15892 LCL were obtained from Coriell. MUTU I and MUTU III were obtained from Jeff Sample and Alan Rickinson. P493-6 LCL were a gift from Elliot Kieff. All B-cell lines were cultured in RPMI-1640 (Invitrogen) supplemented with 10% fetal bovine serum (FBS). 293T cells were cultured in DMEM with 10% FBS. All cell lines were incubated with 1% penicillin-streptomycin (Gibco) in a humidified incubator at 37□C and 5% CO2. All cells were routinely confirmed to be mycoplasma-negative by Lonza MycoAlert assay (Lonza). P493-6 LCLs[122] cultured in the presence of 4-hydroxy tamoxifen (4-HT) and doxycycline (1µg/mL) have EBNA2-driven latency III. The conditional EBNA2-HT fusion protein localizes to the nucleus in the presence of 4-HT, but upon 4HT withdrawal, it is sequestered in the cytosol and degraded. Doxycycline suppresses a modified TET-OFF *c-Myc* cassette. For 4-HT/ doxycycline removal, cells were washed five times with incomplete RPMI. The first two washes included 30-minute incubations in incomplete RPMI media. Cells were then seeded with or without 4-HT/doxycycline for 48 hours before analysis.

### Primary B Cell Isolation and Culture

RosetteSep and EasySep negative isolation kits (Stemcell Technologies) were used sequentially to isolate CD19+ B cells by negative selection, with the following modifications made to the manufacturer’s protocols. For RosetteSep, 40 μL antibody mixture was added per mL of blood and before Lymphoprep density medium was underlayed, prior to centrifugation. For EasySep, 10 μL antibody mixture was added per mL of B cells, followed by 15 μL magnetic bead suspension per mL of B cells. After negative selection, the cells were washed twice with 1x PBS, counted, and seeded for EBV infection studies. Cells were cultured in RPMI-1640 (Invitrogen) supplemented with 10% FBS and penicillin-streptomycin in a humidified chamber at 37□C and 5% CO2. Cells were cultured in RPMI-1640 supplemented with 10% FBS and penicillin-streptomycin in a humidified incubator at 37□C and at 5% CO2.

### Antibodies and Reagents

Antibodies against the following proteins were used in this study: JunB (Cell Signaling Technology, #3753), c-Jun (Cell Signaling Biotechnology, #9165), JunD (Cell Signaling Biotechnology, #5000), β-Actin (Biolegend, #664802), LMP1 (Abcam, ab78113), LMP2A (Abcam, ab59028), EBNA2 PE2 (a gift from Fred Wang), DDX1 (Bethyl, A300-521A-M), Myc (Santa Cruz Biotechnology, SC-40), p100/p52 (EMD Millipore, 05-361), TRAF1 (Cell Signaling Biotechnology, #4715S), HA tag antibody (Cell Signaling Technology, # 3724), BIM (Cell Signaling Biotechnology, #2933), CCND2 (Proteintech, 10934-1-AP), EBNA1 (a gift from Jaap Middeldorp), and EBNA3C A10 (hybridoma from Martin Rowe’s group[123]). The following reagents were utilized in this study at the indicated concentration unless otherwise noted: DMSO (Fisher, BPBP231-100, Doxycycline hyclate (Sigma, D9891-1G, 250 ng/mL for pLIX_402 exogenous LMP1 cell lines, 1μM for maintenance of P493-6 LCL), MEGACD40 ligand (Enzo Life Sciences, 50 ng/ml, ALX-522-110[124]), birinapant SMAC mimetic (Selleckchem, 20μM, #S7015[54]), TNFα (R&D Biosystems, 10ng/ml, #210-TA[124]), IKKβ inhibitor VIII (Apex Bio, A3485, 1μM or 5μM [19].

### B95.8 EBV Preparation and B-cell Infection

B95-8 EBV stocks were prepared from B95-8 producer cells as previously described[125] [126]and stored at −80 degrees C. Infectious titer of freshly thawed EBV was determined by primary B-cell transformation assay. Freshly isolated, de-identified, discarded CD19+ peripheral blood B cells were seeded in RPMI1640 with 10% FBS at a concentration of one million cells/mL for infection studies at an EBV multiplicity of infection of 0.1.

### CRISPR/Cas9 mutagenesis or CRISPR activation

B-cell lines with stable Cas9 or dCas9 CRISPRa casettes were established as described previously[46]. sgRNA constructs were generated as previously described[127] using sgRNA sequences from the Broad Institute Avana or Brunello libraries for Cas9 or Calabrese for CRISPRa. CRISPR editing / overexpression was performed as previously described[128]. Briefly, lentiviruses encoding sgRNAs were generated by transient transfection of 293T cells with packaging plasmids pasPAX2 (Addgene, Plasmid #12260) and pCMV-VSV-G (Addgene, Plasmid #8454). Lentiguide-Zeo (Addgene, Plasmid # 160091), pLentiGuide-Puro (Addgene, Plasmid #52963), and pXPR_502 (Addgene, Plasmid # 96923) plasmids. P3HR-1, Daudi, MUTU I, GM12878, and GM13111 cells stably expressing Cas9 were transduced with the lentiviruses and selected with 3 μg/mL puromycin (Thermo Fisher, Cat#A1113803) for two days before replacement with antibiotic-free media. Zeocin selection was performed using 200 μg/mL Zeocin (Thermo Fisher, # R25005). CRISPR editing was confirmed by immunoblotting four days post puromycin selection and seven days post zeocin selection. For double knockouts, seven days after selection of lentiguide-Zeo JunD or CDKN2C transduced cells, a second transduction was performed using plentiguide-Puro JunB sgRNA vector. For CDKN2C CRISPRa experiments, cells were transduced as above using pXPR_502 vector to deliver CRISPRa sgRNA. Cells were selected using puromycin and then re-seeded four days post selection for cell growth comparative analyses. sgRNAs used in this study were constructed using oligos based on the sequences below:

### RNA-seq

GM12878 cells were transduced with either control sgRNA or JunB sgRNA. Ten days post puroymycin selection, Total RNAs were isolated using Qiagen RNeasy kit (Qiagen, #74104) following the manufacturer’s manual. An in-column DNA digestion step was included to remove any residual genomic DNA contamination. To construct RNA-seq libraries, 500 ng total RNA was used for polyA mRNA-selection, using the NEBNext Poly(A) mRNA Magnetic Isolation Module (New England Biolabs, # E7490S), followed by library construction via NEBNext Ultra RNA Library Prep Kit (New England Biolabs, # E7770S). Each experimental treatment was performed in triplicate. Libraries were multi-indexed, pooled and sequenced on an Illumina NovaSeq XPlus sequencer using paired-end 150 bp reads (Illumina).

RNAseq data analysis: sequencing files were trimmed of the adapter sequences using cutadapt (DOI: https://doi.org/10.14806/ej.17.1.200) and raw reads were aligned to hg38 Human genome assembly using STAR aligner[129]. Bam files were then converted to bigWig for each strand using deepTools bamCoverage function (version 3.5.6) (doi: https://doi.org/10.1093/nar/gkw257). featureCounts was used to obtain per-gene reads and the resulting tables were imported into R and DESeq2[130] was used for differential gene expression analysis and ggplot2 for visualization.

## Data Availability

All RNAseq data sets are available in the GEO omnibus under accession number GSE326036.

## JUNB RNA-seq and ChIP-seq integration

For integration of differentially expressed genes with JunB chromatin occupancy, we downloaded the available narrowPeak file from ENCODE[131] and annotated the peaks using annotatePeak function from the R package ChIPseeker (doi:10.1002/cpz1.585, https://onlinelibrary.wiley.com/share/author/GYJGUBYCTRMYJFN2JFZZ?target=10.1002/cpz1.585, doi:10.1093/bioinformatics/btv145). A 1kb window up- and downstream of each gene transcription start site (TSS) was used to infer overlap between JunB peaks and differentially expressed genes.

BigWig track from ENCODE of the same ChIPseq experiment was uploaded to UCSC Genome Browser, together with CTCF, BATF, IRF4 and all histone marks from GM12878 cell line, for visualization purposes. EBNA3A and 3C ChIP-seq tracks were obtained from https://doi.org/10.1073/pnas.1422580112 (EBNA3A) and https://doi.org/10.1073/pnas.1321704111 (EBNA3C).

## Flow cytometry

For 7-AAD (Thermo Fisher, Cat#A1310) viability assays, cells were harvested and washed twice with 1x PBS supplemented with 2% FBS (Gibco). Cells were then incubated with a 1μg/mL 7-AAD solution in 1x PBS / 2% FBS for five minutes at room temperature, protected from light. Cells were then analyzed via flow cytometry. For CFSE labeling and cell proliferation assays, CellTrace CFSE (Invitrogen, Cat#C34554) solution was prepared according to manufacturer’s instructions. 10^7 cells were resuspended and incubated in one mL of CellTrace working solution for ten minutes in a 37°C / 5% CO2 incubator, protected from light, with the cap of the vessel ajar. Five mL of RPMI 1640 with 10% FBS was added to the stained cells protected from light. Cells were incubated at room temperature for five minutes, protected from light, to remove free dye and prevent toxicity. Cells were then pelleted by centrifugation (300g x 5 minutes, room temperature) and resuspended in fresh RPMI with 10% FBS three times before the cells were seeded in complete media for proliferation analysis experiments at a concentration of 300,000 cells/mL. Labeled cells were analyzed by flow cytometry using a BD FACSCalibur instrument or Cytek Northern Lights instrument and analysis was performed with FlowJo V10.

## PI cell cycle analysis

Cells were collected and washed 2x with 1x PBS. Cells were then fixed and permeabilized using BD Fix/Perm kit (BD, Cat# 554714). After fixation and permeabilization, cells were resuspended in a solution consisting of 800μL of 1x BD Perm Wash, 16 uL of RNAse A (Thermo Fisher, Cat# EN0531), and 16 μL of Propidium Iodide (Thermo Fisher, Cat# P3566). Cells were then incubated for 15 minutes at 37c. After PI staining, cells were washed 2x with 1x PBS. Labeled cells were analyzed by flow cytometry using a BD FACSCalibur or Cytek cytometer and analysis was performed with FlowJo V10.

## CellTiter-Glo

CellTiter-Glo viability assay (Promega) was performed according to the manufacturer’s protocol at the indicated time points. A total of 35μL cells in PBS were used per assay according to manufacturer instruction.

## CRISPR-Cas9 CDKN2C / JunB double knockout

Cas9+ GM12878 and GM13111 were transduced with Lentiguide-Zeo plasmids harboring control or CDKN2C sgRNAs. One week after selection with Zeocin, cells were transduced with pLentiguide-Puro plasmids harboring a separate control or JunB sgRNA. Cell numbers two days following puromycin selection (defined as day 0 of the graph) were set to 1. Live cell numbers were quantitated by CellTiter-Glo assay.)

## Software/data Presentation

Statistical analysis was assessed with Student’s t test using GraphPad Prism 7 software, where NS = not significant, p > 0.05; * p < 0.05 ** p < 0.01; *** p < 0.005. and graphs were made using GraphPad Prism 7. Figure 2A and Figure 7 were created using Biorender.

## Supplementary Figure Legends

**Supplementary Figure 1:**
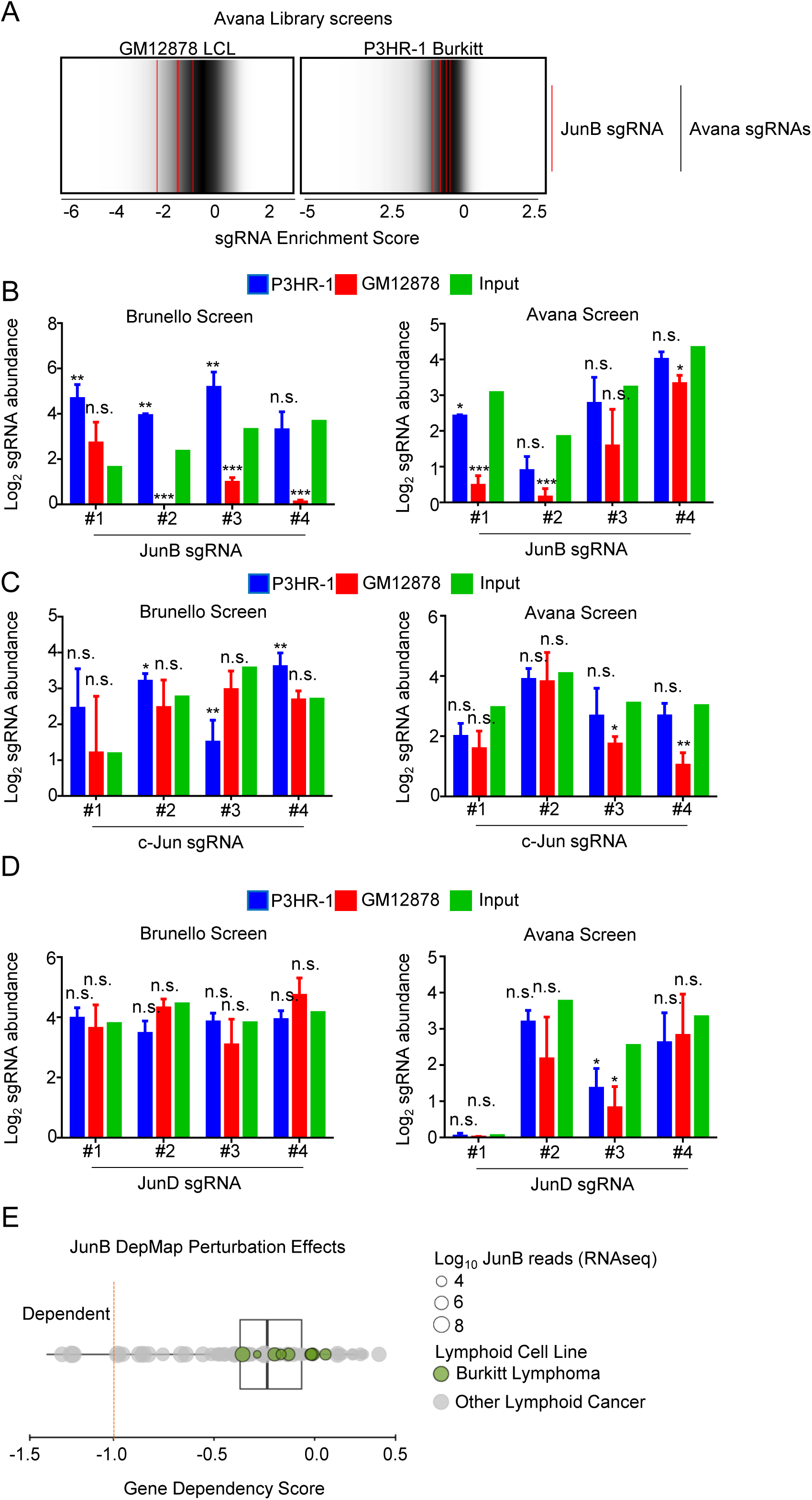
JunB knockout affects lymphoblastoid cell growth. (A) Log2(fold change) of the four guides targeting JunB (red) compared to the distribution of all sgRNA guides from the GM12878 and P3HR-1 Avana dropout screen. Fold change was calculated comparing abundance of sgRNA day 21 post selection versus input. (B) JunB sgRNA abundances in Brunello and Avana dropout screens. Log_2_ JunB sgRNA abundances from GM12878 (red) and P3HR-1 (blue) cells 21 days post library selection were determined via deep sequencing and compared to input library sgRNA abundance (green). (C) c-Jun sgRNA abundances in Brunello and Avana dropout screens. Log_2_ c-Jun sgRNA abundances from GM12878 (red) and P3HR-1 (blue) cells 21 days post library selection were determined via deep sequencing and compared to input library sgRNA abundance (green). (D) JunD sgRNA abundances in Brunello and Avana dropout screens. Log_2_ JunD sgRNA abundances from GM12878 (red) and P3HR-1 (blue) cells 21 days post library selection were determined via deep sequencing and compared to input library sgRNA abundance (green). (E) JunB dependency across CRISPR DepMap lymphoid cell lines (DepMap.org). Gene dependency scores were calculated by comparing JunB sgRNA abundance in individual lymphoid cell lines 21 days post lentivirus library transduction and selection in comparison to input library abundance. Each circle indicates an individual lymphoid cell line. Circle size represents Log JunB reads from RNA-seq analysis (DepMap, Broad, DepMap Public 25Q3 (2025)) Green circles versus clear circles indicate DepMap Burkitt versus non-Burkitt lymphoid cell lines, respectively. P-values were determined by one-sided Fisher’s exact test. * p<0.05, **p<0.005, ***p<0.0005

**Supplementary Figure 2:**
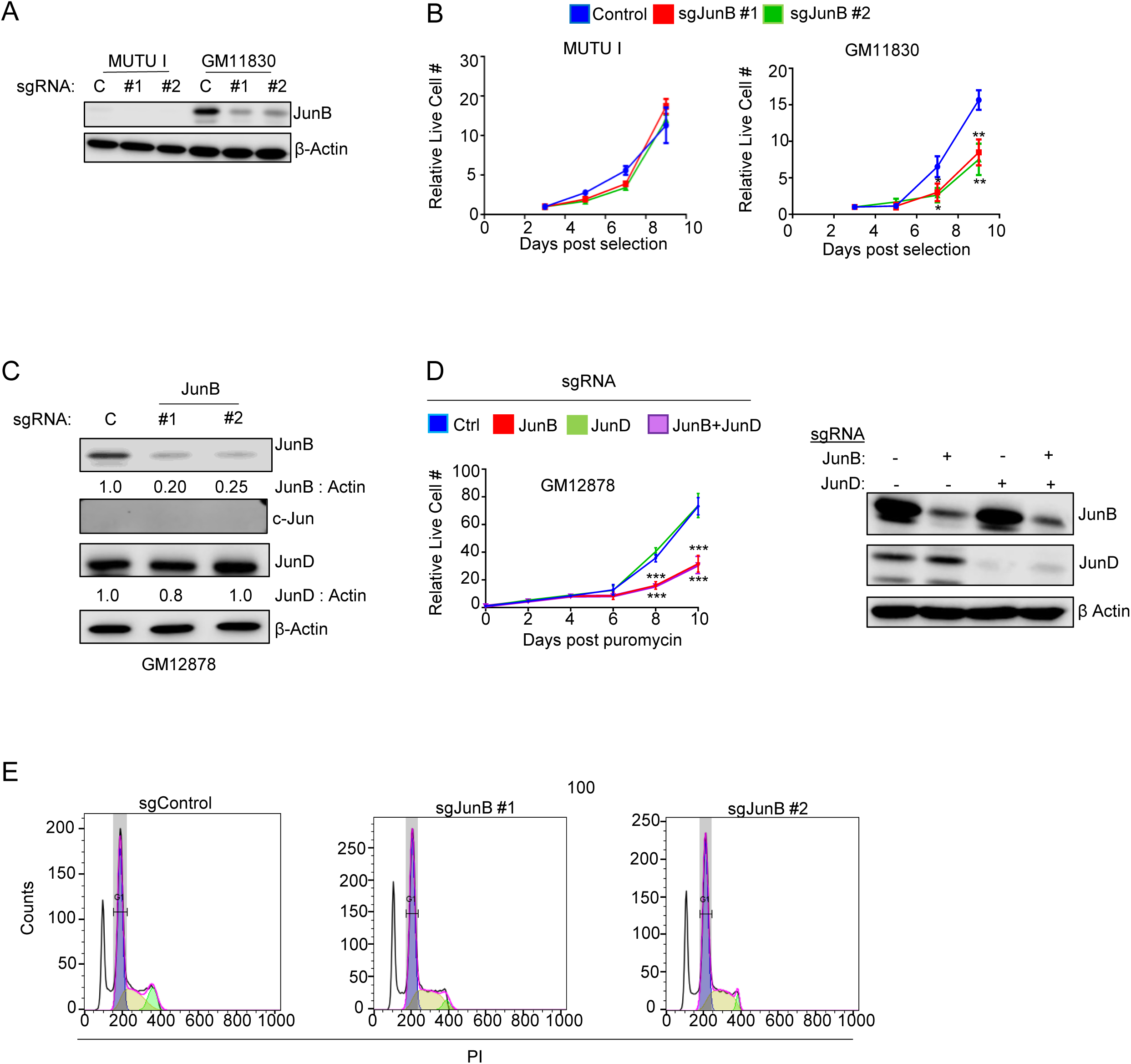
JunB, but not c-Jun or JunD, regulates EBV transformed cell growth and cell cycle. (A) Validation of JunB CRISPR-Cas9 knockout in MUTU I and GM15892 cells. Immunoblot was performed on WCL from Cas9+ MUTU I and GM15892 cells transduced with control or JunB sgRNA seven days post-selection using indicated antibodies. (B) Mean ± SD live cell numbers of Cas9+ MUTU I or GM11830 cells expressing control or JunB targeting single guide RNAs (sgRNA) from n=3 replicates. Cells transduced with lentiviruses expressing the indicated sgRNAs were puromycin selected. Cell numbers two days following puromycin selection (defined as day 0 of the graph) were set to 1. Live cell numbers were quantitated by CellTiter-Glo assay (C) Investigation of impact of JunB on other Jun AP-1 transcription factors. Immunoblot analysis of WCL from Cas9+ GM12878 cells transduced with control or JunB sgRNAs, ten days post selection, with indicated antibodies. Densitometry ratios of JunB and JunD to β-Actin is shown. (D) Investigation of combinatorial impact of JunB and JunD on LCL growth. (Left) Mean ± SD live cell numbers of Cas9+ GM12878 cells expressing control, JunD, JunB, or JunB+JunD targeting single guide RNAs (sgRNA) from n=3 replicates. Cells transduced with lentiviruses expressing control or JunD sgRNA were Zeocin selected two days post transduction. After one week, cells were then transduced with a separate control or JunB sgRNA. Cell numbers two days following puromycin selection (defined as day 0 of the graph) were set to 1. Live cell numbers were quantitated by CellTiter-Glo assay. (Right) Immunoblot validation of JunB and JunD knockouts. WCL of GM12878 cells transduced with both zeocin and puromycin sgRNAs was extracted two days following puromycin selection and immunoblot was performed using indicate antibodies. (E) PI cell cycle stainin of GM12878 cells transduced with JunB sgRNA. Representative FACS cell cycle plots from n=3 replicates of GM12878 cells analyzed ten days post selection. P-values were determined by one-sided Fisher’s exact test. * p<0.05, **p<0.005, ***p<0.0005

**Supplementary Figure 3:**
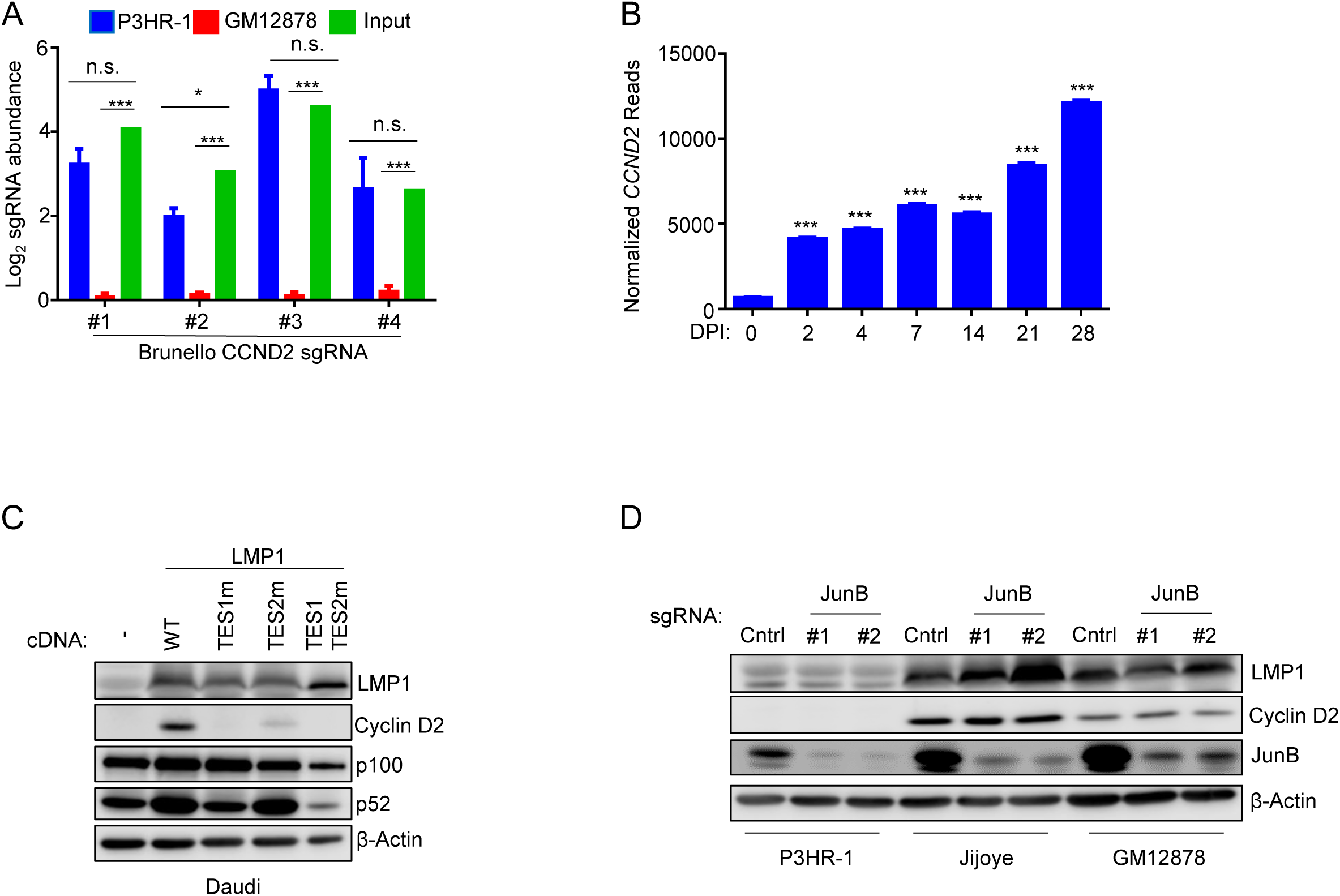
Epstein-Barr virus Latent Membrane Protein 1 regulates CCND2 levels, but not via JunB. (A) CCND2 sgRNA abundances in Brunello P3HR-1 and GM12878 dropout screens. Log_2_ CCND2 sgRNA abundances from GM12878 (red) and P3HR-1 (blue) cells 21 days post library selection were determined via deep sequencing and compared to input library sgRNA abundance (green). (B) CCND2 transcript levels during EBV-mediated B cell transformation. RNAseq was performed on primary B cells infected with B95-8 virus on indicated days post infection (dpi). Normalized reads for *CCND2* are shown. (C) Immunoblot analysis of CCND2 levels in response to LMP1 expression. Daudi Burkitt cells harboring selectively inducible WT or transformation effector site (TES) mutant LMP1 cassettes were mock induced or treated with 250 ng / mL doxycycline for 24 hours. Cells were then harvested and lysates analyzed via immunoblot with indicated antibodies. (D) Effect of JunB knockout on CCND2 levels. Cas9+ P3HR-1, Jijoye, or GM12878 cells expressing control or JunB sgRNAs. Seven days post selection, WCL was extracted from cells and immunoblot performed using indicated antibodies. P-values were determined by one-sided Fisher’s exact test. * p<0.05, **p<0.005, ***p<0.0005

